# Real-time repository-scale spectral search and global molecular networking with HNSW-MS

**DOI:** 10.64898/2026.06.02.729602

**Authors:** Alexander Semenov, Andrew M P Roberts, Elizabeth A Coler, Samarth Gupta, Evguenia Kopylova, Alexey V Melnik, Vladimir Boginski, Alexander A Aksenov

**Author notes:** Contributed equally (Alexander Semenov, Andrew M P Roberts).

## Abstract

Spectral similarity comparison is the basis of mass spectrometry-based metabolomics, underpinning library matching, molecular network construction, and repository searches such as MASST. Public repositories now exceed billion spectra, and exhaustive pairwise comparison is increasingly limiting. We introduce HNSW-MS, which implements Hierarchical Navigable Small World graph indexing natively for mass spectra. It operates directly on GC-MS and LC-MS/MS spectra without embedding, so scores are identical to those of exhaustive comparison and results are fully reproducible. On the benchmark dataset of 8.4 million LC-MS/MS spectra, HNSW-MS is up to 900-fold faster than linear search, recall@1 and recall@10 achieve over 90%, with the recall-speedup tradeoff tunable at query. Such rapid retrieval allows for any query spectrum to be near-instantaneously integrated into a global molecular network. We demonstrate specific examples of use of global networking for molecular discovery, including uncovering a new pathway of stenothricin biosynthesis and a new dipeptide conjugate bile acid.

## Introduction

Mass spectrometry-based metabolomics data analysis rests on the key operation of comparing one fragmentation spectrum to another, such as a reference spectrum from a library. Library matching, molecular networking, and repository search are all built on it^1–6^. Typically, a query spectrum is sequentially compared to each spectrum in a library; until recently, libraries were small enough - GNPS launched in 2016 with 221,083 reference spectra - making exhaustive pairwise comparison tractable, and the design of every major tool reflects that assumption.

This is no longer true, as modern untargeted experiments generate tens to hundreds of thousands of spectra from a single study. Modern instruments such as Thermo Astral or SCIEX ZenoTOF are generating datasets that are orders of magnitude larger than conventional LC-MS platforms. Data in public repositories grow exponentially; GNPS/MassIVE, for example, now exceeds one billion spectra^7,8^. The reference collections have grown alongside, e.g. the GNPS public library now holds over 2.9 million reference spectra. Comparing a query against a library of *N* spectra requires *N* similarity evaluations, each involving peak matching within a mass tolerance window, intensity normalization, and scoring of matched peak pairs. On other hand, network construction requires *N*(*N*-1)/2 comparisons, so that a 50,000-spectrum dataset requires 1.25 billion comparisons. In a practical example, assuming that a batch of ∼100,000 new spectra from a typical untargeted metabolomics experiment needs to be compared against an existing GNPS repository of >1 billion spectra to determine the best match(es) for each new spectrum, the “brute force” linear search would take ∼10^14^ comparisons, which is equivalent to multiple years of computations on a single-core machine. Even in smaller-scale linear search settings, such computations may easily take days to months. Although modern multi-core high performance computers can significantly decrease the computational time, the search is still implemented sequentially, and taking advantage of parallel hardware requires specialized parallel code and additional implementation effort. Acceleration of the comparison operations is of broad value across the field, independent of any particular downstream application, since every workflow that depends on spectral similarity inherits the benefit. Recent developments such as MASST+ have improved search speed by approximately two orders of magnitude through algorithmic optimizations and error-tolerant matching^4^, and Flash Entropy Search has demonstrated real-time matching against combined libraries^9^. In both cases, however, the underlying number of comparisons remains linear with the library size.

Acceleration by orders of magnitude could alter the manner in which public data can be interrogated^3,4^. Conceptually, this would be equivalent to the previous transition from library search to molecular networking within an individual dataset, but extended to the global *repository-scale* data. Under the emerging paradigm of reverse metabolomics, an experimental spectrum is interpreted by placing it in the context of all existing public data rather than within the study that produced it^10,11^. MASST established that a spectrum can be searched against the repository to determine where else the corresponding molecule has been observed. Other tools such as microbeMASST can further attach taxonomic context to that answer^3,10^, while MassQL extends the same logic from whole spectra to spectral patterns^5^. Each of these tools addresses the question of whether a given molecule is present in public data, and in what context. On the other hand, molecular networking addresses a different question, namely which structurally related molecules exist^1,12,13^. With this framing, discovery becomes possible, since a compound that is absent from every reference library may nevertheless be adjacent in a network to compounds that is known^14^. Currently, networking is exclusively conducted on the scale of an individual dataset, and networking on the global scale has not been attempted.

Approximate nearest neighbor (ANN) search offers an alternative route for pairwise spectral similarity comparison. ANNs were originally developed for high-dimensional similarity search in machine learning and information retrieval by reducing the number of distance evaluations by orders of magnitude while returning results close to those of exhaustive search^15,16^. The performance of an ANN algorithm is typically evaluated using metrics that quantify how closely the retrieved results match those obtained by exact nearest-neighbor search, with recall being one of the most commonly used measures^17^. In particular, *Recall@1* is the percentage of queries when the top guess by an ANN algorithm matches the true top match, and *Recall@10* is the proportion of true top-10 matches that are successfully retrieved within the top-10 ranked matches identified by the algorithm. Throughout the manuscript, we will refer to the “performance” or “accuracy” of the search algorithm in the sense of its recall values.

Among the ANN models, Hierarchical Navigable Small World (HNSW) graphs offer a favorable balance of speed, performance, and memory efficiency^16^. However, non-unique input representations and not very high recall values obtained in the respective implementations impede a wide adoption of HNSW in mass spectrometry. The first attempt to apply HNSW to spectral matching^18^ was restricted to EI GC-MS library data and did not operate on raw spectra. The spectra were instead converted into transformed vectors by Word2Vec embedding^18–20^; consequently, while computational speedups were obtained, recall@1 and recall@10 were limited to approximately 37% and 80%, respectively. In that same study, standard HNSW was applied to the Word2Vec embeddings using conventional cosine distance. More consequentially, Word2Vec and other learned embedding methods represent transformations of the original spectra and are not unique, as they depend on the chosen combination of spectral preprocessing and network training parameters, leading to possible irreproducibility. Here we show the implementation of the HNSW method for mass spectrometry (HNSW-MS), to enable real-time repository-scale search and molecular networking.

## Results

The recently introduced Molecular Community Networking (MCN) method^12^ was used to assemble a molecular network of ∼8.4 million molecular features, spanning all public GNPS data^4^. MCN organizes up to 95% of all molecules into connected molecular communities, thereby recovering connections with nodes that conventional networking leaves isolated. HNSW-MS, in combination with MCN, now allows rapid comparisons on the global scale, enabling the transition from simple repository search to integrating any new unknown spectra into the global network in real time.

Specifically, for any query spectrum, HNSW-MS is the first step that is applied to identify its top *k* matches in the repository. This essentially creates the first-order “ego network” of a new unknown spectrum, connecting it to its top k immediate neighbors in the global molecular network. As shown below, HNSW-MS allows one to accomplish this step with high accuracy and in real time. Subsequently, it can be followed by expanding this ego network to add 2nd-, 3rd-, and higher order neighbors. These connections are already known from the global molecular community network topology, which has already been computed on the same GNPS repository dataset^12^ (Figure 1). We show that this combined approach allows one to construct such ego networks for batches of multiple new spectra in real time, and then utilize them to discover new compounds based on the network annotation propagation strategies. Although the presented method is amenable to all types of raw mass spectra, the emphasis of the results presented below is on LC-MS/MS spectra, since there is a global repository available for this type of spectra.

**Figure 1.**
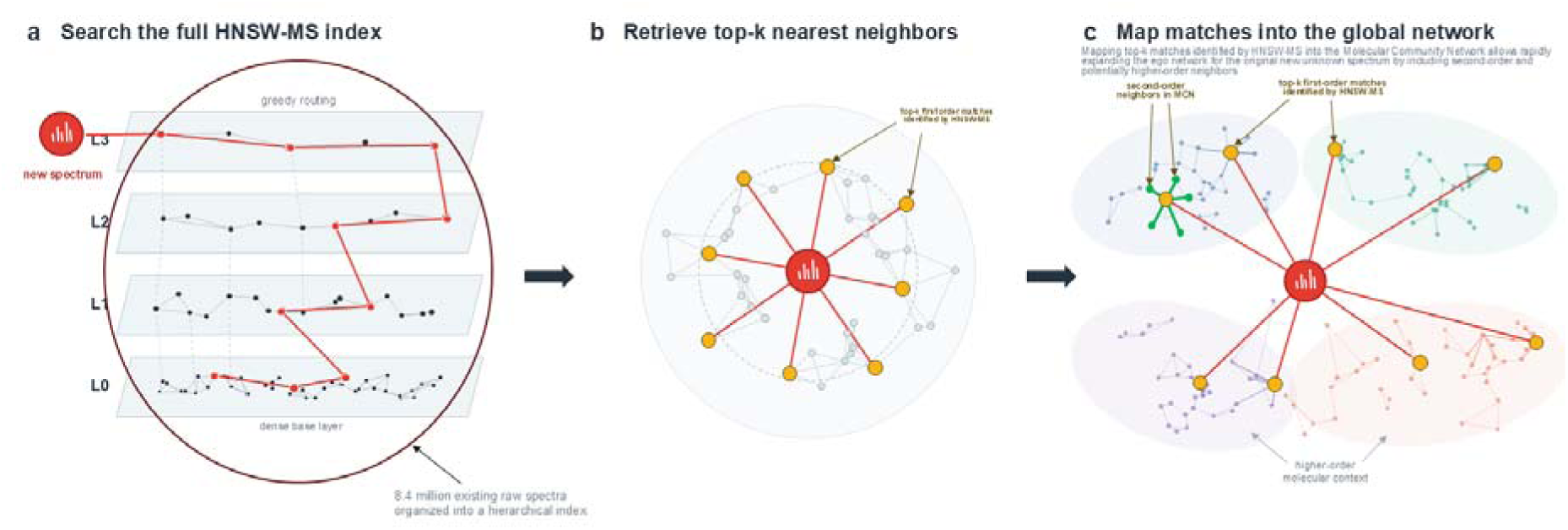
Construction of an ego network for a new spectrum (red node) by: (a) querying a hierarchical graph index of ∼8.4M spectra to (b) retrieve its top-k nearest neighbors using the HNSW-MS approximate nearest-neighbor algorithm (first-order “ego network” construction), then (c) embedding these neighbors into a precomputed global Molecular Community Network so that first-order neighbors immediately expand into higher-order molecular neighborhoods.

### Native indexing of raw mass spectra

A critical distinction of the present implementation is that HNSW-MS operates directly on mass spectral data without requiring transformation into fixed-dimensional vector embeddings, which not only removes the reproducibility problem, but also substantially raises the accuracy.

Each spectrum is represented as a sparse collection of mass-to-charge (*m/z*) values paired with their corresponding intensities. The spectral similarity function performs peak matching within an *m/z* tolerance window (default 0.01 Da), followed by greedy assignment of matched peaks and cosine-based scoring of the matched intensity vectors^1,21,22^. The HNSW-MS preserves the native structure of mass spectral data and ensures that similarity scores computed are identical to those obtained by conventional pairwise comparison, guaranteeing full reproducibility and compatibility with existing spectral analysis pipelines. Importantly, HNSW-MS is agnostic to the scoring function and only affects the number of mutual comparisons; correspondingly, any similarity scoring function can be used. In all the results presented below, the modified cosine similarity was used throughout.

### Library search at repository scale with HNSW-MS

We evaluated HNSW-MS on several datasets of both GC-MS and LC-MS/MS spectra at different scales, between tens of thousands to millions of spectra. Since the focus of this study is on *repository-scale* performance of the proposed method, here we focus on the computational results for the global dataset of 8.4M spectra obtained from all GNPS spectra after clustering to remove redundancy, at the scale of full publicly available repository data^4^. To our knowledge, this is the largest dataset that has been analyzed in this context to date. Performance is reported as speedup over the exact brute force search and the respective Recall@K for K = 1 and 10.

For in-library matching (i.e., there is an exact library match for each queried spectrum) on the global 8.4 million-spectrum dataset, speedups range from 198x to 561x; the fastest setting visits 7,218 nodes on average per query (rather than 8.4 million), and Recall@1 reaches 0.92-0.95 while Recall@10 ranges from 0.74 to 0.89 (Table 1). For the task of out-of-library matching (i.e., queried spectra that are not present in the library) on the same 8.4M dataset, an average of only 6,039 nodes per query were visited in the fastest setting, resulting in over 900x speedup, with Recall@1 and Recall@10 for various settings ranging between 0.79 to 0.92 and 0.8 to 0.93, respectively (Figure 2, Table 2).

**Figure 2.**
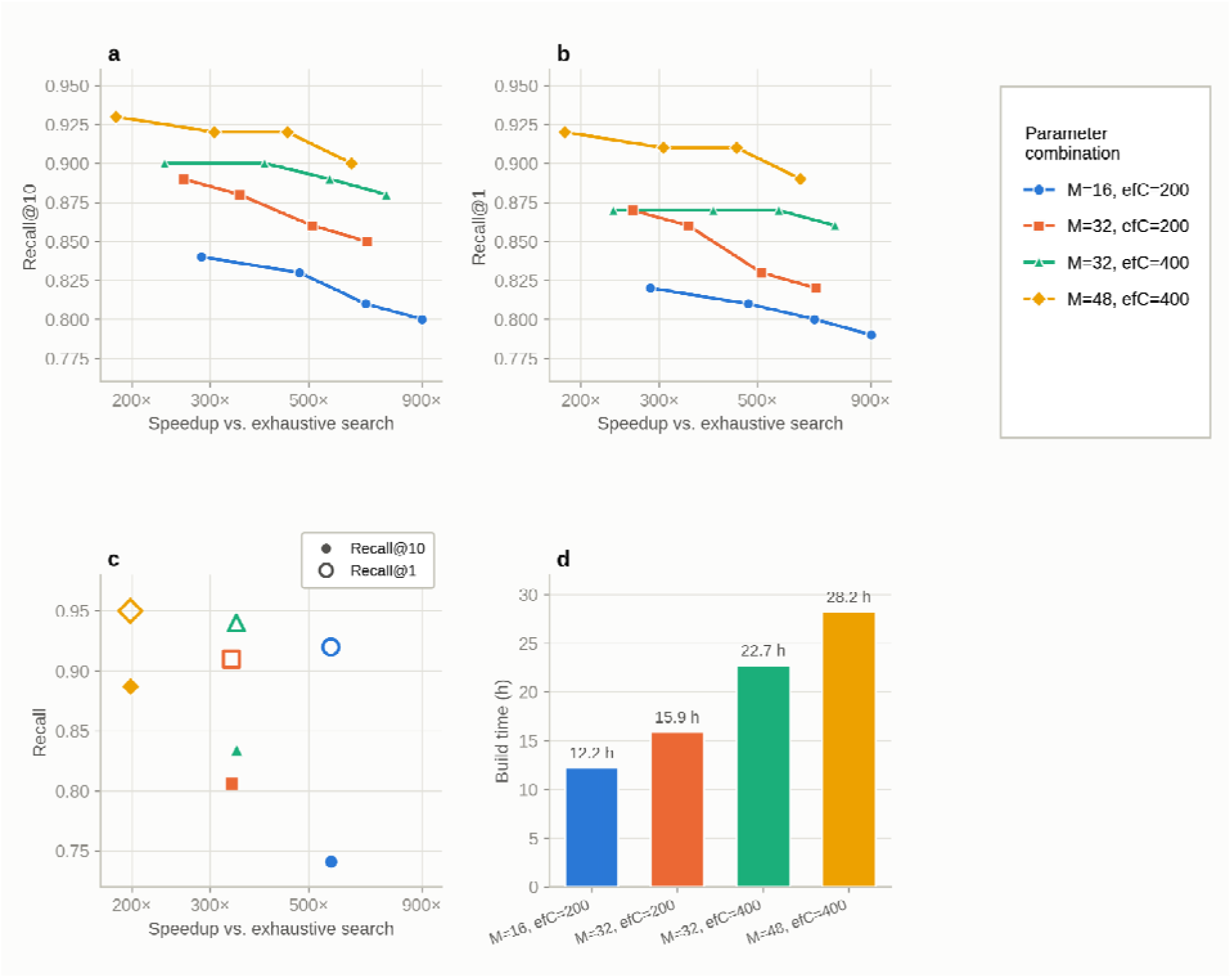
Recall-speed tunability for the global network of ∼8.4 million LC-MS/MS spectra*. a) Recall@10 and b) Recall@1, respectively, as a function of speedup over exhaustive linear search (log scale) for out-of-library matching (i.e. spectra not present in the index). c) Recall for in-library matching achieves higher Recall@1 than out-of-library matching at comparable speedups, reflecting the guaranteed self-match, while Recall@10 is comparable or slightly higher out-of-library. d) One-time index construction (build) time for the same four (M, efConstruction) combinations on the 8.4M-spectrum index; *efSearch* does not affect build time, since it is set at query time on an already-built index. Average nodes visited per query varies inversely with speedup, is omitted for clarity and reported in the source tables in Methods. *n = 100 query spectra per condition (Methods).

**Table 1.** Spectrum matching performance (speedup and recall) on the global dataset of 8.4M LC-MS/MS spectra with different HNSW parameter combinations: in-library matching; n = 100 query spectra per condition.

| $M$ | $efC$ | $efS$ | Speedup | Avg. nodes visited | Recall@10 | Recall@1 |
| --- | --- | --- | --- | --- | --- | --- |
| 16 | 200 | 128 | 561.31 | 7218.35 | 0.741 | 0.92 |
| 32 | 200 | 256 | 335.05 | 12719.5 | 0.806 | 0.91 |
| 32 | 400 | 256 | 343.85 | 11769.7 | 0.834 | 0.94 |
| 48 | 400 | 512 | 198.38 | 20478.7 | 0.887 | 0.95 |

**Table 2.** Spectrum matching performance (speedup and recall) on the global dataset of 8.4M LC-MS/MS spectra with different HNSW parameter combinations: out of library matching; n = 100 query spectra per condition.

| $M$ | $efC$ | $efS$ | Speedup | Avg. nodes visited | Recall@10 | Recall@1 |
| --- | --- | --- | --- | --- | --- | --- |
| 16 | 200 | 128 | 901.25 | 6039 | 0.8 | 0.79 |
| 16 | 200 | 256 | 671.99 | 7826.94 | 0.81 | 0.8 |
| 16 | 200 | 512 | 476.4 | 10647.4 | 0.83 | 0.81 |
| 16 | 200 | 1024 | 286.82 | 15279.7 | 0.84 | 0.82 |
| 32 | 200 | 128 | 676.96 | 7075.52 | 0.85 | 0.82 |
| 32 | 200 | 256 | 509.77 | 9491.45 | 0.86 | 0.83 |
| 32 | 200 | 512 | 349.52 | 13518.3 | 0.88 | 0.86 |
| 32 | 200 | 1024 | 261.31 | 20355.7 | 0.89 | 0.87 |
| 32 | 400 | 128 | 748.05 | 7296.79 | 0.88 | 0.86 |
| 32 | 400 | 256 | 557.85 | 9613.75 | 0.89 | 0.87 |
| 32 | 400 | 512 | 397.55 | 13625 | 0.9 | 0.87 |
| 32 | 400 | 1024 | 236.76 | 20378.7 | 0.9 | 0.87 |
| 48 | 400 | 128 | 624.74 | 8330.72 | 0.9 | 0.89 |
| 48 | 400 | 256 | 448.39 | 11311.4 | 0.92 | 0.91 |
| 48 | 400 | 512 | 306.73 | 16351.8 | 0.92 | 0.91 |
| 48 | 400 | 1024 | 183.98 | 25758.2 | 0.93 | 0.92 |

HNSW-MS was able to achieve high Recall@1 and Recall@10 values for both in-library and out-of-library matching tasks, often exceeding 90% (especially at more “precise” algorithm parameter settings). Recall@1 is generally higher than Recall@10 for in-library matching, whereas the opposite is true for out-of-library matching. This effect may be due to the fact that for in-library matching there is always a guaranteed exact top match for each queried spectrum, and it appears that the algorithm is consistently able to identify that top match, resulting in higher Recall@1 values. On the other hand, for out-of-library matching, there is often no exact match for a queried spectrum within the library, which results in a more “uniform” performance in terms of Recall@1 and Recall@10, with both of these metrics still being substantially high in all algorithm parameter settings (more details in Methods).

Speedup increases approximately logarithmically with dataset size, consistent with the *O*(log *N*) query complexity of HNSW compared to *O*(*N*) for exhaustive search. This scaling behavior indicates that the advantages of HNSW-MS become increasingly pronounced with repository size.

### Tuning recall against speed, and growing the index

Index construction is an incurred one-time cost, scaling approximately as *O*(*N* log *N*) and ranging from 5.6 min (50,000 spectra) for the smallest dataset to 12-28 h for the largest (∼8.4 million spectra) on a single core (Figure 2d, Table 3). Once built, an index is stored for reuse and can be queried indefinitely. The index is also incrementally updatable, with new spectra inserted into the existing graph without reconstruction (Methods, HNSW-MS implementation). This is essential for a continuously growing repository such as GNPS: newly deposited spectra become available to both library search and networking as they arrive, rather than accumulating until a full rebuild is justified. At the highest-accuracy setting the index for the 8.4 million- spectrum collection occupies approximately 10 GB, comparable to the 7 GB of the spectral file it indexes, and both construction and querying were performed within a 30 GB memory allocation (Methods, Computational environment and resource requirements).

**Table 3.** Index construction time (one-time computational cost) for the global dataset of 8.4M LC-MS/MS spectra with different HNSW parameter combinations.

| *M* | *efC* | Build time (min)<br>8.4M entities |
| --- | --- | --- |
| 16 | 200 | 732.46 |
| 32 | 200 | 953.39 |
| 32 | 400 | 1364.08 |
| 48 | 400 | 1693.24 |

The recall-speed trade-off is tunable without rebuilding the index. *M* and *efConstruction*, fixed at construction time, determine a given index’s capacity for accuracy; *efSearch*, set independently at query time, then determines where on that index’s trade-off curve each individual query operates, from maximum acceleration to maximum accuracy (Methods, Parameter selection).

The tolerance for approximation error differs between the two principal use cases. In library search, a false nearest neighbor constitutes a misannotation, so Recall@10 and especially Recall@1 take priority, and higher-recall parameter combinations (those yielding recall above 90% (Tables S1–S3)) are warranted. For network construction, by contrast, the value of the network lies in its topology (the clustering patterns for structurally related spectra into connected components) rather than in the precise identity or rank order of any single edge. Approximate search may substitute edges of near-equivalent similarity without breaking that clustering, so parameters can be tuned for maximum acceleration as the appropriate operating point, not a concession made for networking’s sake. This allows for maximum acceleration for ego network construction for any spectra, to make it a routine, ultra-fast operation feasible at repository scale.

### Integrating new spectra into the global network

Retrieval at the speed enabled by HNSW-MS permits a query mode that was previously either impossible or impractical. A new (unknown) spectrum is submitted to an HNSW-MS index built over the global GNPS molecular network and its *k* nearest neighbors are returned, resulting in a first-order “ego network” of that new spectrum, that is, a subnetwork that contains the new spectrum and its *k* first-order neighbors (*k* was set to 25 as a compromise between neighborhood coverage and the number of nodes requiring manual or AI-assisted triage; densely connected regions may warrant larger *k*). Since these *k* neighbors are in turn already positioned in the global network that already exists (global MCN)^12^, the MCN edges incident to them are traversed outward for *n* hops (*n*=2 here), collecting the annotated and unannotated nodes that surround the query in global chemical space (Fig. 1). Both parameters are adjustable for neighborhood breadth and specificity.

This query returns a mapped molecular neighborhood similar to a conventional molecular network, but for global data. We demonstrate that this global network construction is now a real-time operation that takes milliseconds and can be conducted for a large number of query spectra, readily scalable to entire datasets.

### Practical demonstration of global networking enabled by HNSW-MS

To demonstrate the utility of the global networking approach, selected annotated spectra were used as queries, and their neighborhoods were analyzed by annotation propagation, as described in Methods.

In one example, the global molecular neighborhood of stenothricin G, a cyclic lipodepsipeptide antibiotic from *Streptomyces roseosporus*^23^, contained a homologous series separated by successive CH_2_ increments spanning C48 to C52 acyl compositions, with mass errors within 3.4 ppm and recomputed cosine similarity to the anchor of 0.79-0.91 (Fig. 3), that matched no known spectrum. Fragment ion analysis localized the mass differences to the acyl chain, in that the dominant ions, including the base peak at *m/z* 908, are conserved across the series while only sparse near-precursor ions carry the shift, which is the signature of a variable acyl tail attached to an invariant peptide core (Figure 3).

**Figure 3.**
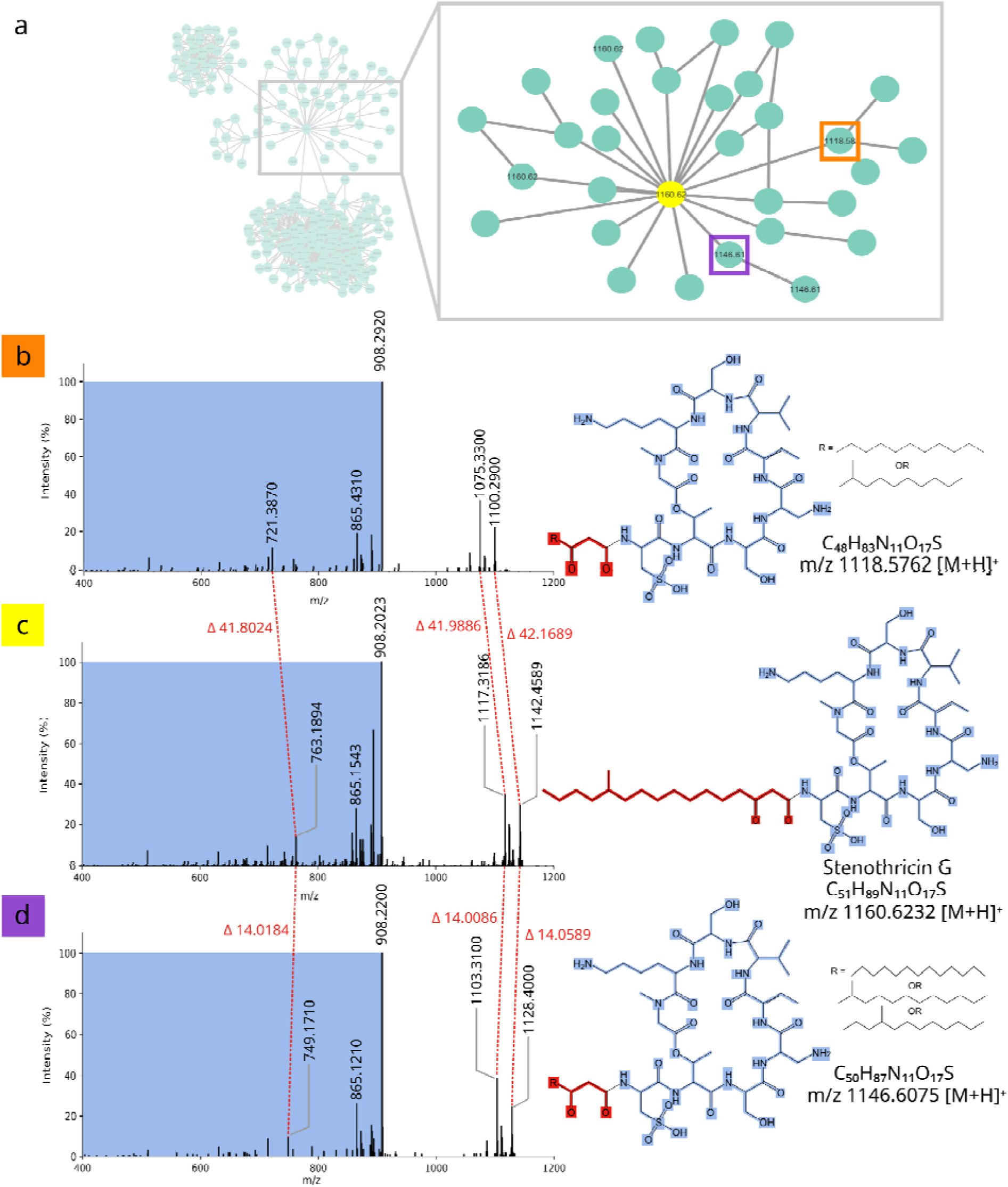
Annotation propagation-based discovery with global molecular networking. a) A molecular neighborhood of the global network of stenothricin G, a cyclic lipodepsipeptide antibiotic from *Streptomyces roseosporus*. b), c), d) MS/MS of the stenothricin G (c) and two postulated variants; the pattern is indicative of the common cyclic peptide core but varying length of the lipid chain. Biosynthetically plausible structures are listed in the Supplemental Table S6.

Both congeners appear novel and carry even-numbered acyl chains of C14 and C16. Branched-chain acyl starter units in gram-positive bacteria derive from branched-chain amino acid catabolism under a strict mapping, in which 2-methylbutyryl-CoA from isoleucine yields odd-chain anteiso fatty acids, isovaleryl-CoA from leucine yields odd-chain iso, and isobutyryl-CoA from valine yields even-chain iso^23–25^. Every currently known stenothricin carries an odd, anteiso-branched chain, as do the daptomycin-family lipopeptides, which are produced by the same strain and are known to be acylated with anteiso-undecanoyl, iso-dodecanoyl and anteiso-tridecanoyl groups^26,27^. Since an even-carbon chain cannot be assembled by the isoleucine route, its presence indicates that the stenothricin acyl-CoA ligase accepts a valine-derived or acetyl-CoA primer, and that the starter unit specificity of this pathway is broader than the known congeners imply. Branch position is not resolvable by CID, since all isomers of a given congener are isobaric. Proposed structures and the biosynthetically admissible branch positions, from which anteiso branching is excluded on carbon-parity grounds, are given in Supplementary Note Table S6.

In another example, the neighborhood of phenylalanine-conjugated cholic acid contained a node at *m/z* 613.384, also observed as its ammonium adduct at *m/z* 630.410, that was matched to cholic acid conjugated with the dipeptide Gly-Phe (calculated [M+H]+ 613.3847, C35H52N2O7; Fig. S1). The CA-Gly-Phe assignment is supported by fragment ions that require an internal amide bond and cannot arise from any single proteinogenic residue, together with the diagnostic trihydroxy-cholanoyl core ions (Methods, Supplementary Note).

Bile acids are conjugated with single amino acids by host enzymes and gut bacteria across a wide range of residues^28–30^, whereas dipeptide conjugation is almost unrecorded. The first example, an N-acyl glycyltaurine conjugate of deoxycholic acid, was reported in rabbit bile in 1998 by authors who examined bile from several hundred species without finding another and concluded that such compounds rarely occur in nature^31^. The second, Met-Glu-cholic acid, emerged from molecular community networking in microbial cultures^12^. CA-Gly-Phe is the third, and three independent observations, made in different systems and by different means, suggest that dipeptide conjugation may be an underdescribed conjugation class rather than an anomaly of rabbit bile. The apparent rarity is likely reflecting how bile acids have been searched, since a conjugate absent from reference libraries is invisible to library matching and a single-dataset network rarely contains the relatives required to recognize it. These examples demonstrate annotation propagation-driven discovery now accessible in global data, enabled by fast and efficient *repository-scale* networking.

## Discussion

We demonstrate how HNSW-MS could resolve the computational bottleneck of spectral similarity search at repository scale without the trade-off that has accompanied previous applications of approximate search in this domain. Because the index is constructed over actual spectra rather than over learned embeddings, the similarity scores it returns are those of the underlying scoring function itself. Additionally, the input format is fixed by the data rather than by a training procedure, so a result obtained with one implementation is reproducible with another. The earlier embedding-based implementation sacrificed these properties in exchange for speed and nonetheless recovered the correct top match in less than half of queries^18^, whereas the present implementation attains speed and accuracy simultaneously, accelerating searches over 8.4 million spectra 200- to 900-fold relative to exact linear scan while returning the best-matching spectrum for over 80% at maximum acceleration and in “accuracy-tuned” settings over 90% of queries.

The practical value of this acceleration scales with the number of searches performed rather than with the duration of any individual one, since a single exhaustive query against 8.4 million spectra is tolerable in isolation. However, this is no longer true if the same query is repeated for every feature of an untargeted dataset, and repeated again each time the repository is updated. Because efSearch parameter is specified at query time and new spectra are accommodated by incremental insertion, mapping of an entire dataset against the whole repository such as GNPS, and its subsequent re-mapping as the repository grows, become routine operations rather than a dedicated computing projects.

Networks are tolerant to non-perfect recall, as networks with slightly altered topology should remain interpretable, while network construction poses greatest computational demands as it scales as ∼N^2^ for the number of pairwise comparisons. Correspondingly, molecular networking construction at repository scale is the application in which the HNSW-MS approach is ostensibly most consequential. Scaling molecular networking -based annotation propagation to all of metabolome, rather than within-dataset, should enable increase in molecular discovery, as any molecule of interest can be placed in the context of its global molecular neighborhood. Both compounds reported here were obtained from two queries directed at spectra that had already been publicly deposited, using a global network that had already been constructed, and no additional data were acquired (Fig. 2). What had been lacking was a means of retrieving, for an arbitrary spectrum, the set of publicly observed structurally related compounds on a timescale compatible with interactive analysis. The “dark matter of metabolomics”, that currently comprises ∼93% of detectable MS features, likely contains both artefactual (in-source fragments, ionoforms), but also truly novel compounds^14^. The finding that the even-carbon stenothricins were present in a producer strain extract, distinguished from the named congeners by nothing more than a CH_2_ count for which no prior hypothesis called, indicates how much remains unexamined in the public repositories.

The principal liability of operating at the global scale becomes interpretive rather than computational. Molecular networking rests on the premise that spectral similarity serves as a proxy for structural similarity. This premise generally holds within an individual dataset, but it begins to fail at a rate that is proportional to probability per each comparison, so that increasing the number of candidate comparisons by six orders of magnitude for the repository scale inevitably increases the number of spurious edges. Both neighborhoods examined here contained such edges. For example, a cluster of nine nodes adjacent to stenothricin G traced entirely to a viral proteomics dataset; moreover, several library annotations within the same neighborhood, assert relationships that their own measured mass differences contradict. The problem is most severe where the spectra are less informative, since low-entropy spectra, i.e. those in which few fragment ions carry most of the intensity, match promiscuously^32^. Entire compound classes may fragment in this manner, lipids prominent among them. Entropy-aware similarity measures would address part of this at the level of scoring^32^ and dense connectivity supported by high-quality MS/MS could reduce it further, but neither eliminates it. Correspondingly, at global scale, the substantial number of connections returned for a given query would be structurally uninformative, and the first task in interpreting a neighborhood is to reject them rather than to explain them.

For this reason, the automated reasoning provided by modern agentic AI tools becomes a requirement rather than a convenience^33^. A neighborhood comprising several hundred nodes, each carrying a mass difference, an annotation of uncertain provenance, along with a fragmentation pattern, cannot be efficiently triaged by manual inspection at the rate at which retrieval produces such neighborhoods, and performing this triage for network of every feature in a dataset is altogether infeasible. The task, which consists in determining whether fragment-level evidence supports a structural relationship or merely a coincidental spectral resemblance, is one of pattern recognition under noise, and language models have recently been shown capable of examining MS/MS spectra and reasoning over fragment evidence in precisely this manner^34–36^. The appropriate division of labor is therefore for an automated agent to discard the unrelated majority and rank what remains with surviving candidates verified by an analyst. The analyses presented here were carried out by the reasoning of this kind; automation of this step remains the principal obstacle between the method and the scale at which it can now be applied.

Several limitations apply. HNSW-MS is approximate and does not guarantee recovery of the exact top-*k* set, particularly under aggressive acceleration settings. Recall@10 declines modestly as collection size grows at fixed parameters, such that recovering it at repository scale requires larger *M* and efSearch with corresponding increases in memory and query time. The current implementation is single-threaded and therefore represents a conservative performance floor; it was left sequential deliberately, since parallelization is an engineering task separable from construction of the index itself, and single-core figures establish what is attainable on commodity hardware, without specialized parallel architecture. A further constraint is that the index and the indexed collection are held in memory in their entirety, which at repository scale amounts to tens of gigabytes (Methods); memory-mapped and compressed storage were not implemented in this version and would reduce this requirement. Parallelization, GPU acceleration, and integration with orthogonal approaches such as MASST+ would yield multiplicative additional gains^4,37–39^. The structures proposed here rest on fragment-level evidence and biosynthetic reasoning and await confirmation against synthetic standards. The additional information such as retention time for chromatography, collision cross section for ion mobility, instrumental settings, library matching score for nodes across network etc. could further augment agentic AI reasoning for more efficient discovery. A strategic query for individual discovery scenarios could further improve performance. Finally, what size network constitutes a well-chosen neighborhood remains an open question that can be addressed empirically going forward, because the cost of each query has become negligible.

## Online Methods

### HNSW-MS algorithm description

The Hierarchical Navigable Small World (HNSW) algorithmaccelerates nearest-neighbor search by constructing, in a preprocessing step, an index in the form of a proximity graph over the reference set^16,40^. In HNSW-MS, each mass spectrum is represented as a node, and edges connect mutually similar spectra, so that similar entries are directly linked and can be reached from one another in a small number of steps. This index is built once and reused for all subsequent queries, replacing exhaustive comparison against every entry with efficient traversal of the graph. The graph is organized hierarchically into layers: a sparse upper layer with few nodes and long-range connections supports rapid coarse localization, while progressively denser lower layers, the lowest containing all entries, support fine-grained local search.

The index is constructed incrementally by inserting entries one at a time. Each new node is assigned a maximum layer at random, and it is connected into the graph by searching the existing structure from the top layer downward to locate its nearest neighbors in each layer it occupies; bidirectional edges are then added between the new node and those neighbors. Retrieval proceeds by the same top-down traversal: beginning at an entry point in the top layer, the search greedily moves to the neighbor most similar to the query, descending layer by layer until it reaches the base layer, where it explores the local neighborhood to collect the final set of nearest candidates. In this way, both construction and search rely on the same greedy graph- navigation procedure.

Three parameters govern the algorithm. *M* is the number of neighbors retained per node and therefore controls the density of the graph; larger *M* produces more edges, improving reachability and accuracy at the cost of a larger index and more comparisons. *efConstruction* (efC) is the size of the candidate list maintained while searching for a new node’s neighbors during index construction; larger *efC* yields a higher-quality graph, with better-chosen connections, at the cost of longer build time. *efSearch* (efS) is the analogous candidate-list size used at query time; it sets the breadth of the search, with larger *efS* examining more nodes and thereby increasing recall at the cost of slower queries. Together, *M* and *efConstruction* determine the quality of the index once built (different “versions” of the index can be built using different combinations of *M* and *efConstruction*, as was the case in this study), while *efSearch* tunes the speed–accuracy trade-off of each individual query.

### HNSW-MS implementation

Application to MS data structure requires extending the standard HNSW algorithm to accommodate arbitrary distance functions, including the non-metric spectral similarity measure used in mass spectrometry. Spectral similarity does not satisfy the triangle inequality^41^ (i.e. knowing that spectrum A to be similar to B, and B to C, places no bound on the similarity of A and C), unlike the common scenario of nearest neighbor algorithms exploiting the triangle inequality to prune candidates during construction. Consequently, during index construction, each node is connected to its nearest neighbors, but standard HNSW then applies pruning: a heuristic that discards some of these candidate edges to cap the number of connections per node and keep the graph sparse. In metric spaces this pruning removes redundant edges without harming search, but under a non-metric spectral similarity it can eliminate connections that the greedy walk actually needs, lowering recall. We therefore relax pruning and allow nodes to accumulate connections more freely, which empirically improves recall in non-metric spaces at a modest cost in memory. Further possible enhancements include precomputed intensity norms, timestamp-based visited arrays, and move semantics to minimize copying during construction.

HNSW-MS implements the hierarchical navigable small world algorithm^16^ in C++ with a spectral similarity function operating directly on raw peak lists. Each spectrum is stored as a sparse vector of *m/z* and intensity pairs. In this implementation, similarity is computed as a modified cosine score: peaks are matched between two spectra within a configurable *m/z* tolerance (default 0.01 Da), matched peak pairs are assigned greedily, and the cosine of the matched intensity vectors is returned. No preprocessing, binning, or embedding is applied, so similarity scores are identical to those produced by conventional pairwise comparison, thus ensuring full reproducibility on all types of mass spectra.

HNSW uses a pre-constructed index. Index construction inserts spectra iteratively. Each spectrum is assigned a maximum layer drawn from a discretized exponential distribution parameterized by *M*, the target number of bidirectional edges per node. A node assigned level L is present in layers 0 through L, so that the exponential decay places most nodes in the base layer alone while a fraction of order 1/M reaches each successive layer; this thinning produces the sparse upper layers used for coarse routing and the dense base layer holding all spectra. Insertion performs a greedy descent from the graph entry point and establishes bidirectional edges to the *M* nearest candidates at each layer, with candidate breadth controlled by *efConstruction*. Implementation optimizations include precomputed intensity norms, timestamp-based visited arrays that avoid per-query clearing, and move semantics during construction. Incremental insertion follows the same procedure as construction: a new spectrum is assigned a layer, greedily routed to its neighborhood, and linked bidirectionally, leaving the remainder of the graph unchanged. Indices therefore grow with the repository without reconstruction.

### Parameter selection

The three HNSW-MS parameters govern the speed vs. accuracy trade-off: higher values increase recall at the cost of search speed, though even high-recall settings remain substantially faster than exact linear search. *M* and *efConstruction* are fixed when the index is built and together set that index’s accuracy ceiling; *efSearch* is specified independently for each query, so a single constructed index can be queried anywhere along its speed-accuracy curve simply by varying *efSearch*, without rebuilding.

*M* controls graph connectivity: higher values create denser graphs with more paths to each node, improving recall at the cost of memory and slightly longer queries (*M*=32 is a reasonable default; *M*=48 for maximum accuracy requirements). *efConstruction* controls search breadth during insertion (*efC*=200 for standard use; *efC*=400 for higher accuracy). *efSearch* controls search breadth at query time and has the most direct effect on the speed/accuracy trade-off - this is the parameter to tune search at query time. For interactive search, *efSearch*=128-256 gives high accuracy at high speed; for batch work where recall is critical, *efSearch*=512-1024 approaches exhaustive accuracy (Table 2). For ego network construction, where edge rank order is not consequential, low *efSearch* settings are appropriate and yield the largest speedups.

### Computational environment and resource requirements

All benchmarks were executed on single cores of nodes of a compute cluster, the nodes of which are equipped with AMD EPYC processors; jobs were dispatched to whichever node was available, so the exact processor model differs between runs, and no node-specific or architecture-specific optimization was applied. The implementation is sequential, therefore the reported timings correspond to single-core throughput and are independent of the core count of the host machine, and, since no specialized parallel architecture is required, comparable performance is expected on commodity hardware. Memory was allocated in excess of requirement, approximately 30 GB per job, so that the spectral collection and the index both resided in RAM throughout construction and querying, and the reported timings therefore do not reflect disk caching effects. For the largest collection, the 8.4 million spectra occupy approximately 7 GB as an MGF file and the corresponding index, at the densest setting used here (*M*=48, *efC*=400), approximately 10 GB on disk; the index is loaded in its entirety at query time, so that peak memory is set by the sum of the two rather than by either alone. Index size grows approximately linearly with the number of spectra; the stored index comprises the peak lists themselves (independent of *M*) plus the graph adjacency, which scales with *M*. Peak lists are stored as uncompressed double precision in this implementation, so these figures reflect the current storage format rather than a requirement intrinsic to the method.

### Benchmark datasets and evaluation protocol

Three collections varying in size were used for library search: 50,000 electron ionization GC-MS spectra from the NIST reference library^42^; 591,719 LC-MS/MS spectra (random subset of GNPS)^13^; and 8.4 million repository LC-MS/MS spectra obtained by clustering all GNPS spectra to remove redundancy^4^. The results for repository-scale search are reported in Tables 1-3 and Figure 2, whereas the results for smaller-scale datasets are given in the Supplementary Note (Tables S1-S5).

For in-library evaluation, 100 query spectra were drawn at random from the indexed collection itself, so that each query has an exact counterpart in the index. Ground truth nearest neighbors were computed by brute-force search using the identical C++ spectral similarity implementation used by HNSW-MS, ensuring that any difference in results arises solely from the approximate graph traversal and not from scoring discrepancies. For each query, the top-*k* library spectra by similarity were retained. Recall@k was computed against this exact top-*k* set (all evaluations used *k*=10 and *k*=1). In particular, recall@1 measures whether the true nearest neighbor is returned as top-1. The exhaustive baseline and the corresponding HNSW-MS queries were timed within the same job on the same node, so that reported speedups are internal ratios and are unaffected by node-to-node variation.

We further evaluated HNSW-MS in an out-of-library matching setting, querying 100 external spectra (not present in the repository) against each library. On the 591,719-spectrum library subset, recall@10 ranged from 0.69 to 0.97 and rose monotonically with *M, efC*, and *efS*, while speedup over exhaustive search fell from ∼401× to ∼28×; on the 8.4M LC-MS/MS library, recall@10 reached 0.80-0.93 with speedups of ∼900× down to ∼184×, the larger speedups reflecting the much slower linear baseline at repository scale (Figure 2b, c). As expected, higher search effort (larger *efS*, and thus more nodes visited) traded throughput for accuracy, and denser graphs (higher *M*) shifted the whole curve upward.

Compared with the in-library experiments at matched parameters and query count, out-of-library recall@10 is somewhat higher (at *M*=48, *efC*=400, *efS*=512: 0.92 versus 0.887 on the 8.4M collection and 0.95 versus 0.90 on the 591,719-spectrum collection), whereas exact top-1 agreement is higher in-library (0.95 versus 0.91 on the 8.4M collection). An in-library query has a guaranteed identical self-match at the top of the ranking, which the index recovers consistently; an external query’s nearest neighbour is merely the closest available reference, so top-1 placement is harder while the composition of the top-10 set is less sensitive to it. Overall, out-of-library retrieval remains accurate and one to three orders of magnitude faster than exhaustive search, with *M*48/*efC*400 offering the best accuracy and *M*16/*efC*200 the best speed.

### Global molecular network and community edges

The global molecular network over public GNPS spectra (∼8.4M spectra) was constructed previously using MASST+ and organized into molecular communities by Molecular Community Networking^4,12^. It is used here as an existing resource and was not rebuilt. Molecular community network edges connect nodes that conventional cosine networking leaves unconnected, including nodes assigned to different communities in a single-dataset network. An HNSW-MS index was built over the spectra of this network (Figure 2d).

### Query time for ego network construction

A query spectrum is submitted to the HNSW-MS index and its *k* nearest neighbors by modified cosine similarity are retrieved (*k*=25 throughout this work). These seed nodes define entry points into the global network. From the seeds, MCN edges are traversed for *n* hops (*n*=2 throughout this work), and the union of nodes reached is returned as the molecular neighborhood of the query, with node precursor masses, library annotations where present, community assignments, and edge similarity scores. Both *k* and *n* are user-specified. Retrieval and traversal complete in milliseconds on a single core. A single query against the 8.4 million- spectrum index requires approximately 5.6 ms, compared to approximately 3.1 s by exhaustive search, so that ten thousand queries (typical number of features in an untargeted LC-MS/MS dataset) can be processed in under 1 min rather than in roughly 9 hours.

### Annotation propagation

For each neighbor, the sign of the mass difference was resolved against the node table and a candidate formula fitted within 5 ppm. Spectral similarity to the anchor was then recalculated independently from the raw peak lists (modified cosine, 0.5 Da tolerance) rather than taken from the network’s precomputed edge scores. Fragment ions were classified as core ions (matching the anchor exactly) or substituent-retaining ions (matching only after the precursor mass shift is added back); a candidate with mostly unshifted core ions and shifted ions confined to the near- precursor region indicates the mass difference is localized to a single substituent.

Stenothricin congeners were assigned by holding the literature-defined peptide core fixed^26,43^ and subtracting its formula from each precursor mass to isolate the acyl portion. This procedure reproduces the masses of the known congeners stenothricin C and H to within a few ppm. Structures were built with RDKit and every proposed formula and stereocenter validated computationally. Acyl branch positions were constrained by the reported branch positions of stenothricins A, C and G and by the parity rules of branched-chain fatty acid biosynthesis^24,44^, and are presented as a biosynthetic hypothesis, since CID does not distinguish saturated methyl branch isomers.

For the bile acid neighborhood, an exhaustive combinatorial search was performed over the 210 dipeptide mass combinations of the 20 proteinogenic amino acids, conjugated by amide bond to seven bile acid cores with permitted water-loss states capped by the hydroxyl count of each core, and extended to 1,540 tripeptide combinations including ornithine, citrulline and GABA for unresolved high-mass nodes. Mass matches were accepted only with supporting fragment evidence. CA-Gly-Phe (*m/z* 613.384 [M+H]+, calculated 613.3847, Δ 0.7 mDa; *m/z* 630.410 [M+NH4]+) is supported by the phenylalanine immonium ion at *m/z* 120.079, the y1 ion at *m/z* 166.085 corresponding to protonated free phenylalanine and placing it at the C-terminus, the y2 ion at *m/z* 223.107 corresponding to release of the intact Gly-Phe unit, a sequential water-loss ladder from *m/z* 412.281, and the trihydroxy-cholanoyl core ions at *m/z* 337.251 and 319.239 (Fig. S1). The y1 and y2 ions are peptide bond cleavage products and require an internal amide bond; no single proteinogenic residue is isomeric with Gly-Phe.

### AI-assisted annotation and triage

The initial pass for the annotation propagation reported here was carried out through interactive reasoning with a large language model (Claude Opus 4.8, Anthropic, San Francisco, CA), employed as an aid under human supervision, rather than an autonomous agent. The model was prompted to resolve signed mass differences to candidate molecular formulae within the stated ppm tolerance, to classify the fragment ions of each candidate as core ions or substituent-retaining ions relative to the anchor spectrum, to enumerate and screen the dipeptide and tripeptide combinations described above against the observed fragmentation, and to reason from the rules of branched-chain fatty acid biosynthesis to the admissible acyl starter units, within the neighborhood of corresponding MCN. The combinatorial enumeration itself, together with all formula and structure validation, was subsequently executed in code (RDKit). Every assignment that was retained was then verified by the analyst against the raw spectra.

## Supplemental Materials

HNSW_Nat_Supplemental

## Data and code availability

HNSW-MS source code is available at https://github.com/Alexander0/HNSW-MS. Benchmark spectral collections are available from GNPS^1^ and NIST^42^. Spectra, network exports, annotated MS/MS assignments, and SMILES for the proposed structures are provided in Supplementary Data. Prebuilt indices are not distributed owing to their size; the parameters required to reproduce them are given in Table 3.

## Supporting information

HNSW_Supplemental

## Acknowledgments

A.A. was supported by Agilent and IEATLIGHT. A.S. and V.B. were supported by Agilent.

## Author contributions

A.A. conceptualized the methodology for MS-based metabolomics. A.S. and V.B. devised the mathematical underpinning of the HNSW-MS. A.S. performed code engineering. E.K, A.M. evaluated and beta-tested the code and workflow. S.G. developed localized code implementation. A.A., A.R., E.C. examined data for the manuscript. A.A. provided supervision for the project. A.A., A.R., V.B., and A.S. wrote and edited the manuscript. All authors reviewed and approved the manuscript.

## Declaration of conflict of interests

A.A.A. is a co-founder of Arome Science, Inc. (https://www.arome-science.com/) and BileOmix, Inc. (bileomix.com/). A.A.A. is also a founder of GreenScent, Inc. All other authors declare no conflict.

## Declaration of generative AI and AI-assisted technologies in the manuscript preparation process

During the preparation of this work, the authors used Claude (Anthropic, model Opus 4.8) to propose candidate structures via annotation propagation. Claude was also used to assist with paraphrasing and refining scientific language. After using this tool/service, the authors reviewed and edited the content as needed and take full responsibility for the content of the article.

## Notes

### Competing Interest Statement

AAA is a cofounder of Arome Science, BileOmix, and GreenScent Inc.

### Summary of Updates

Figures 1 & 2 revised; Added Figure 3; author affiliations updated; supplementary files created; all tables were edited, some moved to supplementary for clarity.

