## Supplementary material for "Real-time repository-scale spectral search and global molecular networking with HNSW-MS": HNSW_Supplemental

### **Supplemental Materials**

**Additional Computational Results for HNSW-MS on Smaller-Scale Datasets**

| **M** | **efC** | **efS** | **Speedup** | **Avg. nodes visited** | **Recall@10** | **Recall@1** |
| --- | --- | --- | --- | --- | --- | --- |
| 16 | 200 | 64 | 53.75 | 663.98 | 0.95 | 0.96 |
| 16 | 200 | 512 | 25.62 | 1935.54 | 1.00 | 1.00 |
| 32 | 200 | 64 | 45.18 | 828.34 | 0.96 | 0.94 |
| 32 | 200 | 512 | 18.65 | 2863.48 | 1.00 | 1.00 |

**Table S1.** Spectrum matching performance (speedup and recall) on a benchmark dataset of 50,000 GC-EI spectra with different HNSW parameter combinations: in-library matching

| **M** | **efC** | **efS** | **Speedup** | **Avg. nodes visited** | **Recall@10** | **Recall@1** |
| --- | --- | --- | --- | --- | --- | --- |
| 16 | 200 | 128 | 104.90 | 1913.85 | 0.79 | 0.85 |
| 32 | 200 | 256 | 50.93 | 4370.23 | 0.85 | 0.88 |
| 32 | 400 | 256 | 51.45 | 4470.20 | 0.87 | 0.85 |
| 48 | 400 | 512 | 21.45 | 12522.10 | 0.90 | 0.94 |

**Table S2.** Spectrum matching performance (speedup and recall) on the dataset of 591,719 LC-MS/MS spectra (subset of global GNPS dataset) with different HNSW parameter combinations: in-library matching

| **M** | **efC** | **efS** | **Speedup** | **Avg. nodes visited** | **Recall@10** | **Recall@1** |
| --- | --- | --- | --- | --- | --- | --- |
| 16 | 200 | 128 | 401.21 | 1058.18 | 0.69 | 0.65 |
| 16 | 200 | 256 | 228.82 | 1722.87 | 0.76 | 0.72 |
| 16 | 200 | 512 | 118.7 | 3217.43 | 0.85 | 0.81 |
| 16 | 200 | 1024 | 72.52 | 5823.26 | 0.88 | 0.82 |
| 32 | 200 | 128 | 275.25 | 1655.56 | 0.78 | 0.71 |
| 32 | 200 | 256 | 163.73 | 2891.59 | 0.85 | 0.8 |
| 32 | 200 | 512 | 87.98 | 5542.14 | 0.91 | 0.82 |
| 32 | 200 | 1024 | 46.4 | 10302.9 | 0.94 | 0.88 |
| 32 | 400 | 128 | 295.83 | 1646.99 | 0.82 | 0.8 |
| 32 | 400 | 256 | 161.97 | 2896.03 | 0.84 | 0.82 |
| 32 | 400 | 512 | 84.02 | 5678.94 | 0.94 | 0.88 |
| 32 | 400 | 1024 | 43.19 | 10575.9 | 0.96 | 0.92 |
| 48 | 400 | 128 | 187.32 | 2404.7 | 0.86 | 0.83 |
| 48 | 400 | 256 | 108.45 | 4294.21 | 0.91 | 0.84 |
| 48 | 400 | 512 | 54.88 | 8187.23 | 0.95 | 0.87 |
| 48 | 400 | 1024 | 27.62 | 15421 | 0.97 | 0.87 |

**Table S3.** Spectrum matching performance (speedup and recall) on the dataset of 591,719 LC-MS/MS spectra (subset of global GNPS dataset) with different HNSW parameter combinations: out of library matching

| **M** | **efC** | **Build time (s) 50K** |
| --- | --- | --- |
| 8 | 100 | 334.3 |
| 8 | 200 | 563.1 |
| 16 | 100 | 368.6 |
| 16 | 200 | 646.6 |
| 32 | 100 | 480.5 |
| 32 | 200 | 862.1 |

**Table S4.** Index construction time for a benchmark dataset of 50,000 GC-EI spectra

| **M** | **efC** | **Build time (min)** 591K entities | **Build time (min)** 8.4M entities |
| --- | --- | --- | --- |
| 16 | 200 | 37.65 | 732.46 |
| 32 | 200 | 57.83 | 953.39 |
| 32 | 400 | 105.7 | 1364.08 |
| 48 | 400 | 174.34 | 1693.24 |

**Table S5.** Index construction time for larger benchmark datasets

**Proposed Structures of Novel Branch-Chain Acyls**

| **m/z** | **Molecular Formula** | **SMILES** | **Proposed Structures** |
| --- | --- | --- | --- |
| 1118.5762 | C_48_H_83_N_11_O_17_S | CCCCCCCCCCCC(CC(NC(C(NC(C(NC(C(NC(C(NC(C(NC(C(NC(C(NC(C(N(C1)C)=O)CCCCN)=O)CO)=O)C(C)C)=O)=CC)=O)CN)=O)CO)=O)C(OC1=O)C)=O)CS(=O)(O)=O)=O)=O | 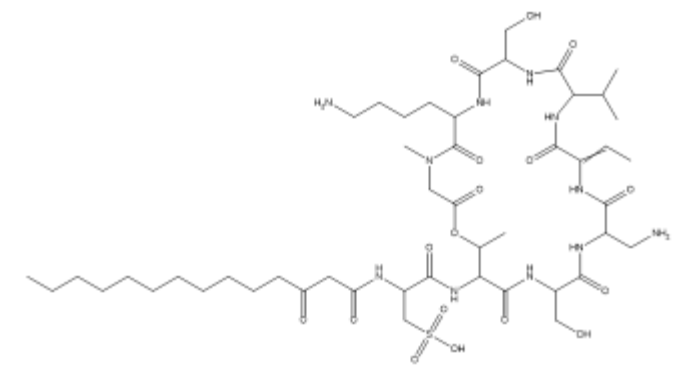 |
| 1118.5762 | C_48_H_83_N_11_O_17_S | CCCCCCCC(C)CC(CC(NC(C(NC(C(NC(C(NC(C(NC(C(NC(C(NC(C(NC(C(N(C1)C)=O)CCCCN)=O)CO)=O)C(C)C)=O)=CC)=O)CN)=O)CO)=O)C(OC1=O)C)=O)CS(=O)(O)=O)=O)=O | 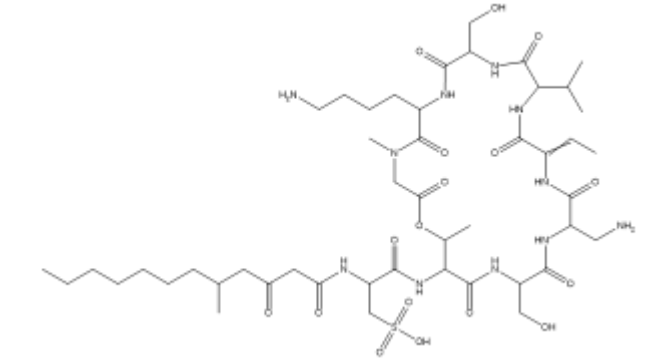 |
| 1146.6075 | C_50_H_87_N_11_O_17_S | O=C(CC(NC(C(NC(C(NC(C(NC(C(NC(C(NC(C(NC(C(NC(C(N(C1)C)=O)CCCCN)=O)CO)=O)C(C)C)=O)=CC)=O)CN)=O)CO)=O)C(OC1=O)C)=O)CS(=O)(O)=O)=O)CCCCCCCCCCCCC | 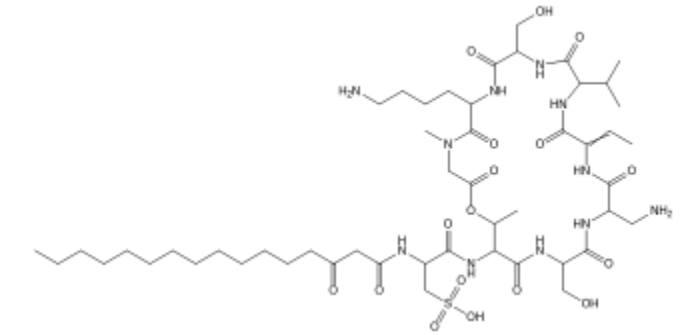 |
| 1146.6075 | C_50_H_87_N_11_O_17_S | O=C(CC(NC(C(NC(C(NC(C(NC(C(NC(C(NC(C(NC(C(NC(C(N(C1)C)=O)CCCCN)=O)CO)=O)C(C)C)=O)=CC)=O)CN)=O)CO)=O)C(OC1=O)C)=O)CS(=O)(O)=O)=O)CC(C)CCCCCCCCCC | 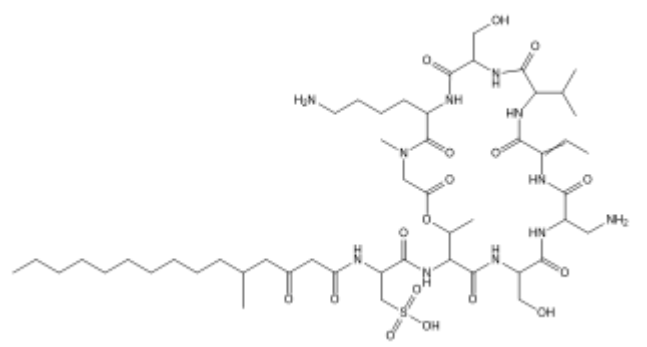 |
| 1146.6075 | C_50_H_87_N_11_O_17_S | CCCCCCCCC(CCCC(CC(NC(C(NC(C(NC(C(NC(C(NC(C(NC(C(NC(C(NC(C(N(C1)C)=O)CCCCN)=O)CO)=O)C(C)C)=O)=CC)=O)CN)=O)CO)=O)C(OC1=O)C)=O)CS(=O)(O)=O)=O)=O)C | 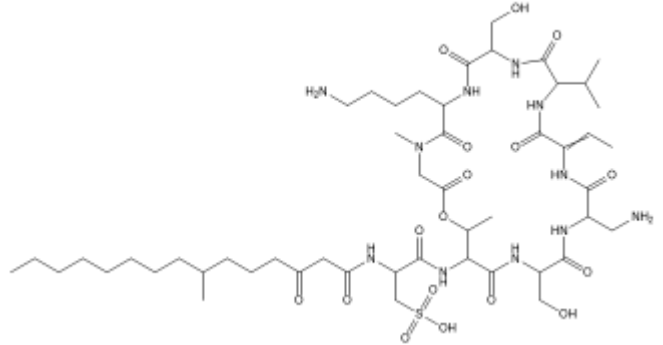 |

**Table S6.** Additional possible Stenothricin structures from the Stenothricin MCN.

**CA-Gly-Phe Assignment**

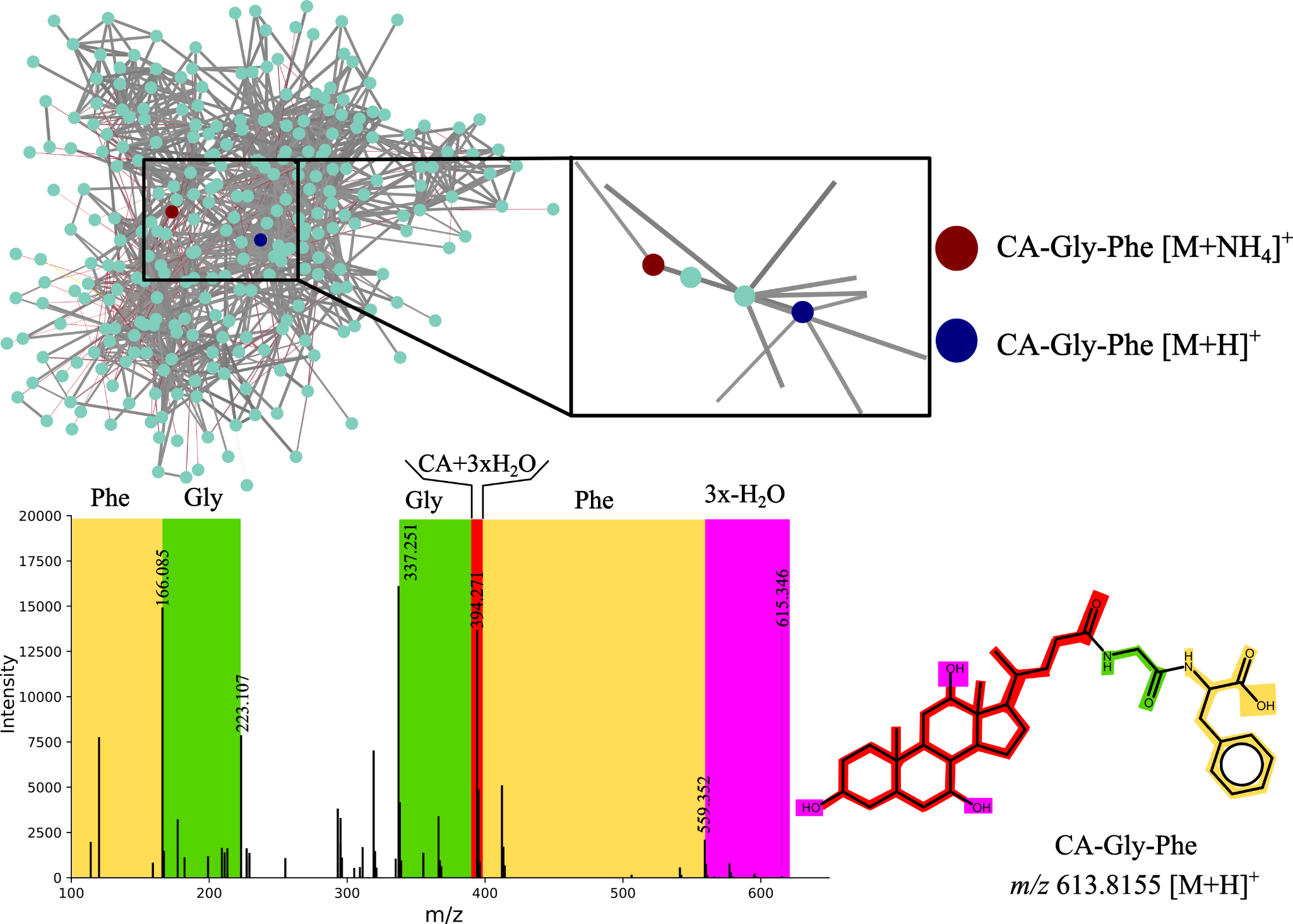

**Figure S1.** Bile acid connections in the molecular network and the proposed structure for unannotated nodes highlighted within the community with m/z 630.410 (red) and 613.384 (blue).
